# Vesicle-mediated transport of ALIX and ESCRT-III to the intercellular bridge during cytokinesis

**DOI:** 10.1101/2022.11.22.517322

**Authors:** Sascha Pust, Andreas Brech, Catherine Sem Wegner, Harald Stenmark, Kaisa Haglund

**Affiliations:** Department of Molecular Cell Biology, Institute for Cancer Research, Oslo University Hospital, Montebello, N-0379 Oslo, Norway; Centre for Cancer Cell Reprogramming, Institute of Clinical Medicine, Faculty of Medicine, University of Oslo, Montebello, N-0379 Oslo, Norway

## Abstract

Cellular abscission is the final step of cytokinesis that leads to the physical separation of the two daughter cells. The scaffold protein ALIX and the ESCRT-I protein TSG101 contribute to recruiting ESCRT-III to the midbody, which orchestrates the final membrane scission of the intercellular bridge. Here, we addressed by which mechanisms ALIX and the ESCRT-III subunit CHMP4B are transported to the midbody. Structured illumination microscopy revealed gradual accumulation of ALIX at the midbody, resulting in the formation of spiral-like structures extending from the midbody to the abscission site, which strongly co-localized with CHMP4B. Live-cell microscopy uncovered that ALIX appeared together with CHMP4B in vesicular structures, whose motility was microtubule-dependent. Depletion of ALIX led to structural alterations of the midbody and delayed recruitment of CHMP4B, resulting in delayed abscission. Likewise, depletion of the kinesin-1 motor KIF5B reduced the motility of ALIX-positive vesicles and caused delayed recruitment of ALIX, TSG101 and CHMP4B to the midbody, accompanied by impeded abscission. We propose that ALIX, TSG101 and CHMP4B are associated with endosomal vesicles transported along microtubules by kinesin-1, leading to their directional transport to the cytokinetic bridge and midbody, thereby contributing to their function in abscission.

## Introduction

Cytokinetic abscission, which leads to the separation of two daughter cells, is tightly regulated in time and space and involves the recruitment of a multitude of proteins to the midbody (Echard et al., 2004; Eggert et al., 2004; Fededa and Gerlich, 2012; Fremont and Echard, 2018; Skop et al., 2004). The midbody is a fundamental protein platform in the intercellular bridge and essential for the initiation of abscission (Addi et al., 2018; D’Avino and Capalbo, 2016; Fremont and Echard, 2018; Guizetti et al., 2011). During the last two decades, molecular mechanisms and the spatiotemporal control of cytokinetic abscission have been increasingly elucidated (Eggert et al., 2006; Elia et al., 2013; Fremont and Echard, 2018; Glotzer, 2005; Green et al., 2012; Mierzwa and Gerlich, 2014). A core component of the midbody is the centralspindlin complex, which recruits the centrosomal protein CEP55, followed by the accumulation of the ESCRT-I (endosomal sorting complex required for transport-I) subunit TSG101 and ALIX (Carlton et al., 2008; Carlton and Martin-Serrano, 2007; Christ et al., 2016; Lee et al., 2008; Morita et al., 2007; Zhao et al., 2006). Both proteins in turn coordinately recruit the ESCRT-III component CHMP4B (charged multivesicular body protein 4B) to the midbody by independent mechanisms (Carlton et al., 2008; Carlton and Martin-Serrano, 2007; Christ et al., 2016; Morita et al., 2007). Abscission initiates by ESCRT-III polymerization into helical filaments that spiral and constrict at the abscission site (Addi et al., 2018; Carlton, 2010; Elia et al., 2012; Goliand et al., 2018; Guizetti and Gerlich, 2012; Guizetti et al., 2011; Lafaurie-Janvore et al., 2013; Mierzwa and Gerlich, 2014; Morita et al., 2007). Finalization of abscission involves F-actin depolymerization, microtubule severing and ESCRT-III-driven membrane scission (Addi et al., 2018; Carlton, 2010; Carlton et al., 2008; Connell et al., 2009; Dambournet et al., 2011; Elia et al., 2012; Fremont et al., 2017; Guizetti and Gerlich, 2012; Lafaurie-Janvore et al., 2013; Mierzwa et al., 2017; Morita et al., 2007).

Trafficking of vesicles with different cargo occurs during early and late steps of cytokinesis, and interference with membrane and protein transport impairs the stability of the intercellular bridge and cytokinesis (Fremont and Echard, 2018; Goss and Toomre, 2008; Montagnac et al., 2008; Neto et al., 2011; Sagona et al., 2010; Schiel et al., 2013; Schiel and Prekeris, 2013). Electron microscopy (EM) and live-cell imaging studies have revealed the existence of membrane vesicles in the intercellular bridge (Dambournet et al., 2011; Fielding et al., 2005; Guizetti et al., 2011; Kouranti et al., 2006; Mullins and Biesele, 1977; Neto et al., 2013; Schiel et al., 2011; Schiel et al., 2012). Besides transport of different protein cargo, vesicles are also crucial for membrane insertion (secretory vesicles) and for remodeling of the membrane lipid composition, in particular of phosphoinositides (Atilla-Gokcumen et al., 2014; Fremont and Echard, 2018; Ng et al., 2005). Many Rab GTPases are present at the cleavage furrow and in the intercellular bridge (Fremont and Echard, 2018; Kelly et al., 2009; Kumar et al., 2019; Militello et al., 2013; Pellinen et al., 2008; Schiel and Prekeris, 2013). In terms of directional protein transport during cytokinesis, most functional studies have focused on Rab11- and Rab35-positive vesicles. Both proteins are required for normal furrow ingression and regulate endosomal recycling pathways required for cytokinesis (Chesneau et al., 2012; Hickson et al., 2003; Klinkert and Echard, 2016; Kouranti et al., 2006; Wilson et al., 2005). FIP3- and Rab35-positive endosomes accumulate at the future abscission sites, where FIP3 endosome fusion promotes the formation of secondary ingressions and Rab35 endosomes recruit effectors required for F-actin clearance, thereby ensuring normal ESCRT-III recruitment and abscission (Dambournet et al., 2011; Fremont et al., 2017; Guizetti et al., 2011; Klinkert and Echard, 2016; Schiel et al., 2011; Schiel et al., 2012).

Vesicle transport into the intercellular bridge is mediated by molecular motor proteins (Fremont and Echard, 2018; Gromley et al., 2005). Microtubule-based transport depends on kinesin and dynein motor proteins, with kinesins mediating plus-end- and dyneins responsible for minus-end-directed transport (Zhu et al., 2005). Rab11-FIP3 endosomes are transported along microtubules (Li et al., 2014) and Rab35 endosomes are moving in the bridge (Dambournet et al., 2011), but if this occurs by kinesin-mediated transport is not known. Interestingly, bi-directional movement of Rab11 and Rab35 has been reported in early bridges, whereas in late bridges they are mostly stationary (Dambournet et al., 2011; Montagnac et al., 2009; Simon and Prekeris, 2008; Takahashi et al., 2011). FIP3 can directly interact with Rab11 and the bi-directional movement of FIP3-positive endosomes is dependent on a switch between plus-end and minus-end motor proteins (KIF5B and dynein) and controlled by Arf6 and JIP4 (Montagnac et al., 2009). In MDCK cells the plus-end-directed motor KIF3A/B mediates the transport of Rab11-FIP5-positive endosomes to the center of the bridge (Li et al., 2014). On the other hand, also actin-dependent motor proteins, such as myosin VI (Arden et al., 2007), are involved in cytokinesis, and the transport of Rab8-positive vesicles might depend on myosin VI (Fremont and Echard, 2018). In human cells the motor proteins for Rab35-positive or secretory vesicle transport into the bridge are not known (Fremont and Echard, 2018).

Even though functional roles of ALIX and other midbody-associated proteins, such as CHMP4B and TSG101, during cytokinesis have been studied in detail (Christ et al., 2016; McCullough et al., 2008; Morita et al., 2007), relatively little is known about the mechanisms and spatiotemporal dynamics by which these proteins are transported into the intercellular bridge and eventually to the midbody.

In this study, we address the spatiotemporal dynamics and mechanisms of ALIX and the ESCRT-III subunit CHMP4B transport during cytokinesis. In accordance with Addi et al. (Addi et al., 2020), we demonstrate a gradual accumulation of ALIX, in co-localization with CHMP4B, at the midbody and toward the abscission site in spiral-like structures. Moreover, our data suggest a highly dynamic recruitment of ALIX and CHMP4B to the midbody and that ALIX and CHMP4B-positive vesicles undergo directional transport along microtubules to the periphery of and into the intercellular bridge, eventually contributing to their midbody recruitment. We further demonstrate that the microtubule motor kinesin-1 promotes transport of ALIX-positive vesicles, the recruitment of ALIX, TSG101 and CHMP4B to the midbody as well as accurate timing of cytokinetic abscission.

## Results

### Highly dynamic populations of ALIX and CHMP4B gradually accumulate at the midbody and form spiral-like structures

At late stages of cytokinesis, ALIX is well-documented to be recruited to the midbody and to promote CHMP4B midbody recruitment (Carlton et al., 2008; Christ et al., 2016; Morita et al., 2007). During maturation of the cytokinetic bridge the localization of ALIX and CHMP4B changes from the appearance as two parallel stripes at the midbody to a cone-shaped structure (Addi et al., 2020; Elia et al., 2011; Lafaurie-Janvore et al., 2013). Furthermore, ALIX and CHMP4B show a high degree of colocalization and eventually both proteins are recruited to the abscission site (Addi et al., 2020). We were interested to understand the spatiotemporal dynamics of ALIX recruitment to the midbody in further detail. To do this, we first examined endogenous ALIX at different stages of midbody maturation in HeLa K cells using structured illumination microscopy (SIM) (Fig. 1A). We detected early appearance of endogenous ALIX in dotted structures at the midbody (Fig. 1A, panel I.), followed by the formation of two ALIX-positive ring-like structures adjacent to the midbody (Fig. 1A, panels II.-III.), formation of elongated cone-like ALIX-positive structures extending from the midbody (Fig. 1A, panels IV.-V.) and finally appearance of ALIX at the abscission site (Fig. 1A, panel VI. and Fig. 1B). We also monitored a high degree of association between ALIX and CHMP4B during this process by SIM (Fig. 1C). ESCRT-III-dependent contractile filaments form at the constriction sites in the intercellular bridge (Goliand et al., 2018; Guizetti and Gerlich, 2012; Guizetti et al., 2011). Accordingly, the detailed analysis of our 3D SIM data revealed that the cone-like structures of ALIX and CHMP4B in two-dimensional microscopy correspond to three-dimensional spiral-like structures (Fig. 1C and Video 1A). These findings are consistent with the morphological changes of ALIX and CHMP4B at the midbody described previously (Addi et al., 2020; Lafaurie-Janvore et al., 2013). To investigate the dynamics of ALIX and CHMP4B recruitment to the midbody, we performed FRAP (fluorescence recovery after photobleaching) analysis of ALIX and CHMP4B at the midbody. The FRAP data showed a very fast recovery of both proteins at the midbody, indicating the existence of highly dynamic protein populations of ALIX and CHMP4B (Fig. 1D and Video 1B). On the other hand, no substantial recovery occurred at post-mitotic midbody remnants (Fig. 1D and Video 1C). Thus, our data suggest that ALIX and CHMP4B gradually accumulate at the midbody in a highly dynamic manner, leading to the formation of spiral-like structures and their eventual accumulation at the abscission site.

**Figure 1.**
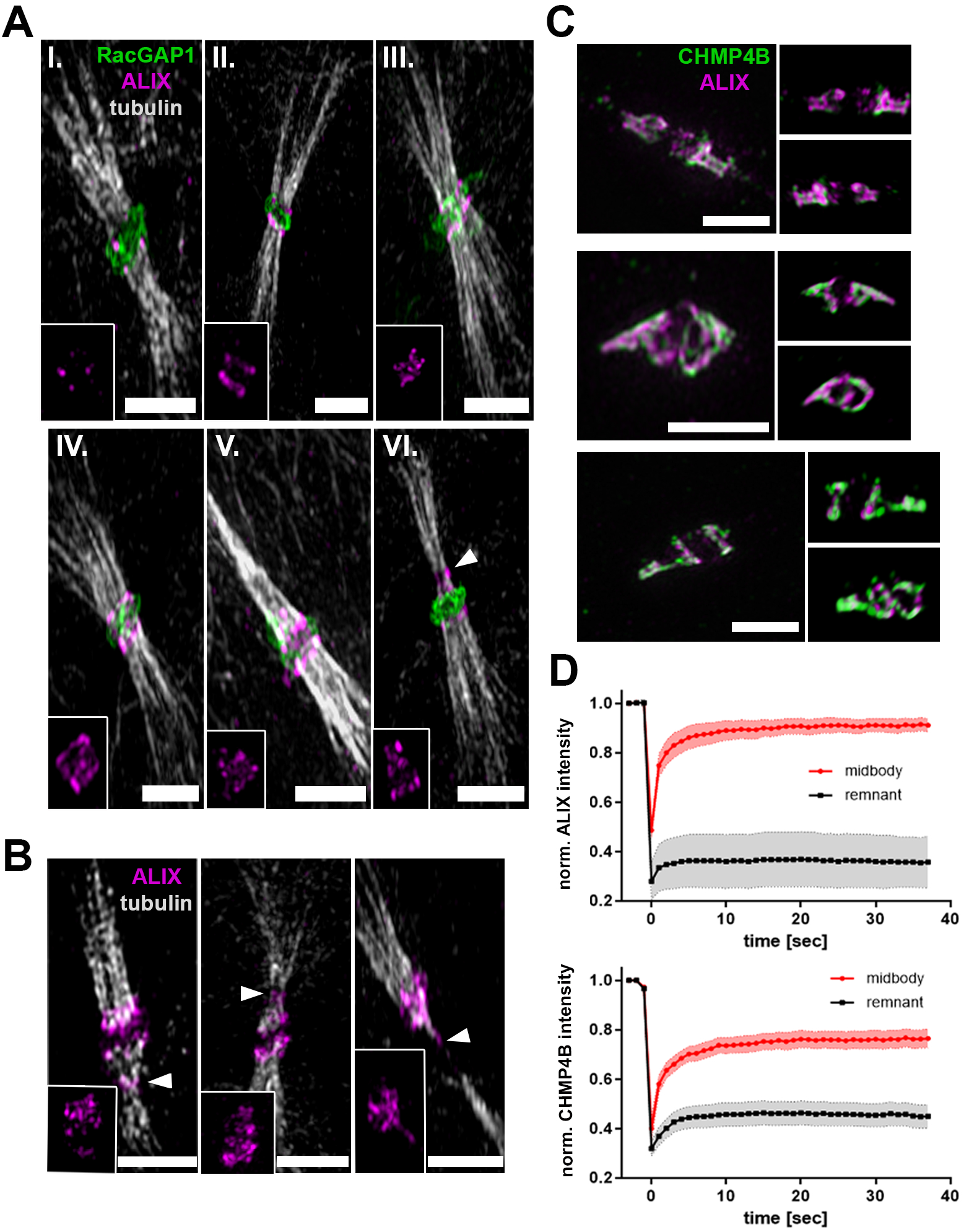
Gradual accumulation of ALIX and CHMP4B at the midbody and colocalization in spiral-like structures. **(A+B)** Sequential accumulation of ALIX at the midbody. 3D SIM microscopy of fixed cells stained for ALIX (magenta), RacGAP1 (green) and tubulin (grey) at different progressive stages of cytokinesis (I.-VI.). **(B)** Recruitment of ALIX to the secondary ingression and abscission sites in fixed cells stained for ALIX (magenta) and tubulin (grey). Inlays show structural changes of ALIX at the midbody, secondary ingression and abscission sites (arrowheads) in (A) and (B). **(C)** Colocalization of ALIX (magenta) and CHMP4B (green) and formation of spiral like structures at the midbody in fixed cells. Representative images show projections of 3D reconstructed SIM data at different visual angles. Cells were stained for ALIX (magenta), CHMP4B (green) and tubulin (grey). Scale bars in (A-C) = 2 μm. See also Video 1A. **(D)** FRAP analysis of ALIX and CHMP4B dynamics at the midbody (red) and in post abscission midbody remnants (grey). Normalized intensities from four independent experiments of cells stably expressing ALIX-mCherry (top) or CHMP4B-GFP (bottom) are plotted (number of FRAP experiments: ALIX-midbody = 12, ALIX-remnant = 11, CHMP4B-midbody = 10, CHMP4B-remnant = 11). See also Video 1B-C.

### ALIX depletion leads to delayed CHMP4B recruitment and abscission timing as well as altered intercellular bridge and midbody morphology

ALIX plays an evolutionarily conserved role in promoting cytokinesis (Carlton et al., 2008; Carlton and Martin-Serrano, 2007; Christ et al., 2016; Eikenes et al., 2015; Morita et al., 2010; Morita et al., 2007) and depletion of ALIX delays the completion of the abscission process (Addi et al., 2020; Eikenes et al., 2015). To characterize the cytokinetic effects of ALIX depletion in our cellular system, HeLa K cells, we performed siRNA-mediated depletion of ALIX in these cells (Fig. 2A). We analyzed abscission time in live cell imaging experiments in control and ALIX-depleted cells by measuring the time starting from the formation of a stable cytokinetic bridge until the microtubules in the bridge were severed (Fig. 2B). Consistent with earlier findings (Addi et al., 2020; Christ et al., 2016), abscission was strongly delayed upon ALIX depletion in HeLa K cells compared to control HeLa K cells (Fig. 2B). ALIX directly interacts with and promotes recruitment of CHMP4B to the midbody (Carlton et al., 2008; Morita et al., 2007). Consistently, we observed substantially delayed recruitment of CHMP4B to the midbody upon ALIX depletion in live cell imaging experiments in HeLa K cells (Fig. 2C and Suppl. Fig. 1). Accompanied with these phenotypes, ALIX depletion also led to the appearance of elongated cytokinetic bridges (Fig. 2D). The analysis of our high-resolution SIM data also revealed that knockdown of ALIX interfered with the structural properties of the midbody (Fig. 2E). Midbodies stained with an antibody against RacGAP1 (Rac GTPase Activating Protein 1), a component of the centralspindlin complex that forms a core structure at the midbody, exhibited significant morphological alterations in ALIX knockdown cells (Fig. 2E). In control cells, the midbody appeared as a ring-shaped structure with of a diameter of approximately 1 μm (Fig. 2E). In ALIX-depleted cells these structures were substantially enlarged and often accompanied with filament-like extensions associated to the midbody structure (Fig. 2E and Videos 2A-C). These structural alterations might be a consequence of the delayed abscission process associated with an erroneous regulation of mechanical forces within the cytokinetic bridge. Altogether, ALIX depletion in HeLa K cells resulted in significantly delayed CHMP4B midbody recruitment and cytokinetic abscission, increase in intercellular bridge length as well as alterations in the morphology of midbodies.

**Figure 2.**
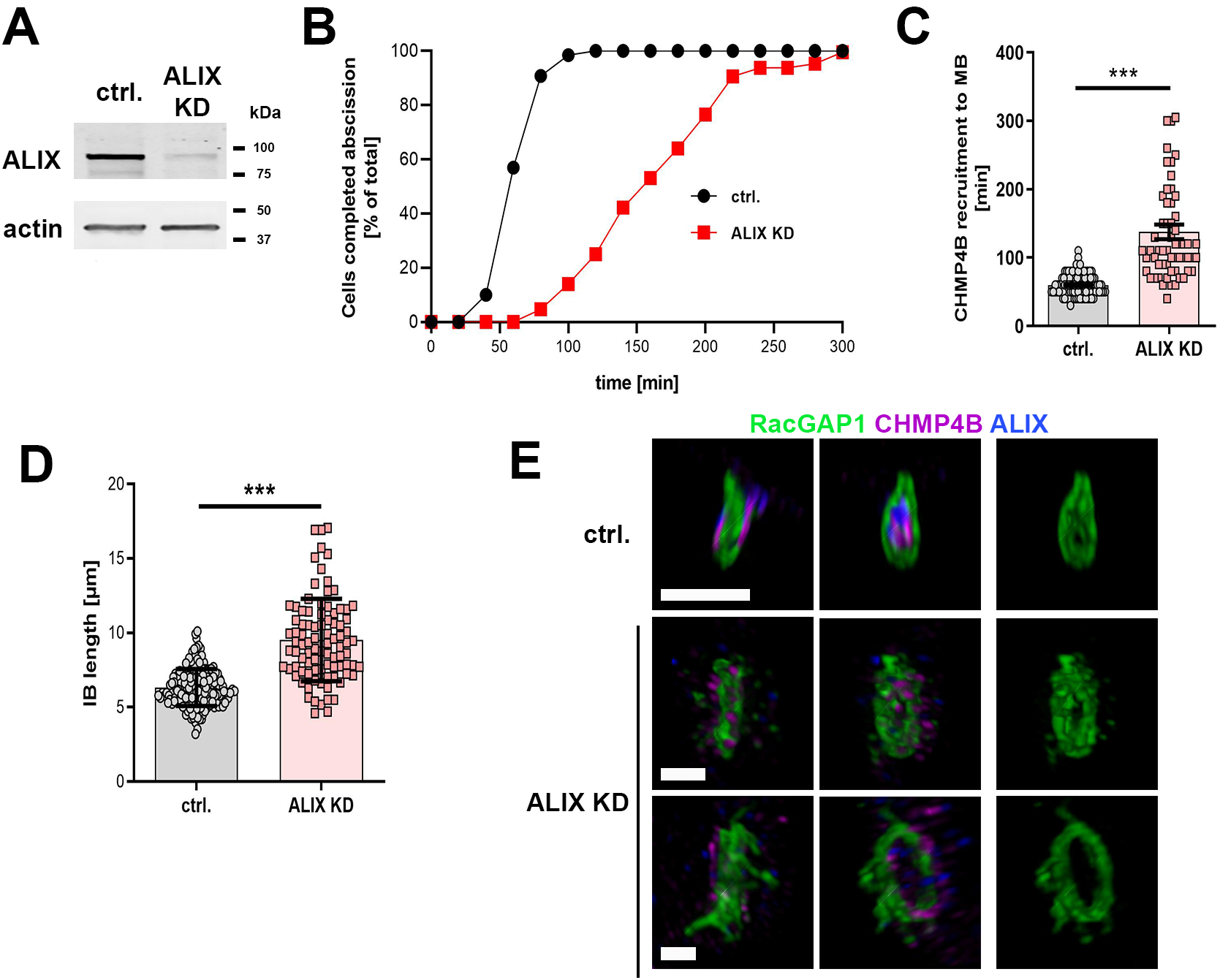
Cytokinetic defects upon ALIX knockdown. **(A)** Western blot showing knockdown (KD) efficiency of siRNA-induced ALIX depletion after 3 days of transfection. **(B)** Cumulative frequency plot showing the time interval between intercellular bridge formation and abscission upon control and ALIX siRNA treatment as indicated (*n* ≥ 60 cells per treatment from three independent experiments; control: 63.8 ± 1.5 min; ALIX KD: 169.2 ± 8.4 min [mean time 50% of cells completed abscission ±SEM]; P < 0.001). **(C)** Scatter plot showing the time interval between bridge formation and appearance of CHMP4B at the midbody (MB) in live cell imaging analysis upon control (ctrl.) and ALIX siRNA (ALIX KD) treatment as indicated (*n* ≥ 60 cells per treatment from three independent experiments; control: 59.9 ± 1.4 min; ALIX KD: 137.8 ± 10.6 min [±SEM]; P < 0.001). **(D)** Quantification of the length of the intercellular bridge (IB) from live cell imaging data (*n* ≥ 100 cells from three independent experiments; control: 6.33 ± 0.1 μm; ALIX KD: 9.5 ± 0.29 μm [±SEM]; P < 0.001). **(E)** Altered morphology of the midbody ring upon ALIX depletion. Projections of reconstructed 3D SIM data at different visual angles are shown. Cells were fixed and stained for RacGAP1 (green), CHMP4B (magenta) and ALIX (blue). Control cells show a compact midbody ring with ALIX and CHMP4B accumulation on both sides of the ring structure. In contrast to control cells, ALIX-depleted cells display enlarged and less compact midbody rings with filamentous extensions. Scale bars = 1 μm.

### ALIX and CHMP4B localize to vesicular structures transported along microtubules

In order to elucidate how ALIX is transported to different cellular destinations, particularly to the cytokinetic bridge and the midbody, we performed live cell imaging of HeLa K cells stably expressing fluorescently labelled ALIX and/or CHMP4B. In earlier studies, ALIX and CHMP4B have been detected in dotted, vesicle-like structures in interphase cells (Katoh et al., 2003). Here, we find that such ALIX-positive vesicular structures are dynamic and transported along the microtubule network in interphase cells (Fig. 3A, upper panel and Video 3A). Accordingly, microtubule depolymerization by nocodazole treatment led to aggregation and immobilization of ALIX-positive structures in interphase cells (Fig. 3A, lower panel and Video 3A). We found that the ALIX-positive vesicles were often also positive for CHMP4B in interphase cells, and consistently, we could detect CHMP4B co-transport with ALIX (Fig. 3B, upper panel). After it became evident that ALIX and CHMP4B localized to vesicle-like structures that are transported along microtubules, we investigated their transport in cells undergoing cytokinesis. As in interphase cells, ALIX and CHMP4B also localized to vesicular structures in cytokinetic cells and both proteins were detected to be co-transported over long distances to the periphery of the cytokinetic bridge (Fig.3B, lower panel and Video 3B). The existence of endosomal vesicles in the cytokinetic bridge has been previously documented and their crucial role in protein and membrane transport during cytokinesis is well established (Baluska et al., 2006; Fielding et al., 2005; Schiel et al., 2011; Schiel et al., 2012; Wilson, 2005). To test whether the intracellular accumulations of ALIX and CHMP4B were associated to vesicular structures we treated HeLa K cells stably expressing fluorescently labelled ALIX and CHMP4B cells with CellBrite^®^ Steady 650 membrane dye. We observed in live cell imaging co-localization and co-transport of ALIX and CHMP4B together with CellBrite-positive vesicles in interphase cells (Video 3C) as well as to the periphery of the intercellular bridge in cytokinetic cells (Video 3D). Co-localization of ALIX and CHMP4B in the intercellular bridge was also confirmed by super-resolution imaging (Fig. 3C). Thus, in interphase and cytokinetic cells, ALIX colocalizes with CHMP4B on vesicular structures and both proteins are transported on such structures in a directed and microtubule-dependent manner.

**Figure 3.**
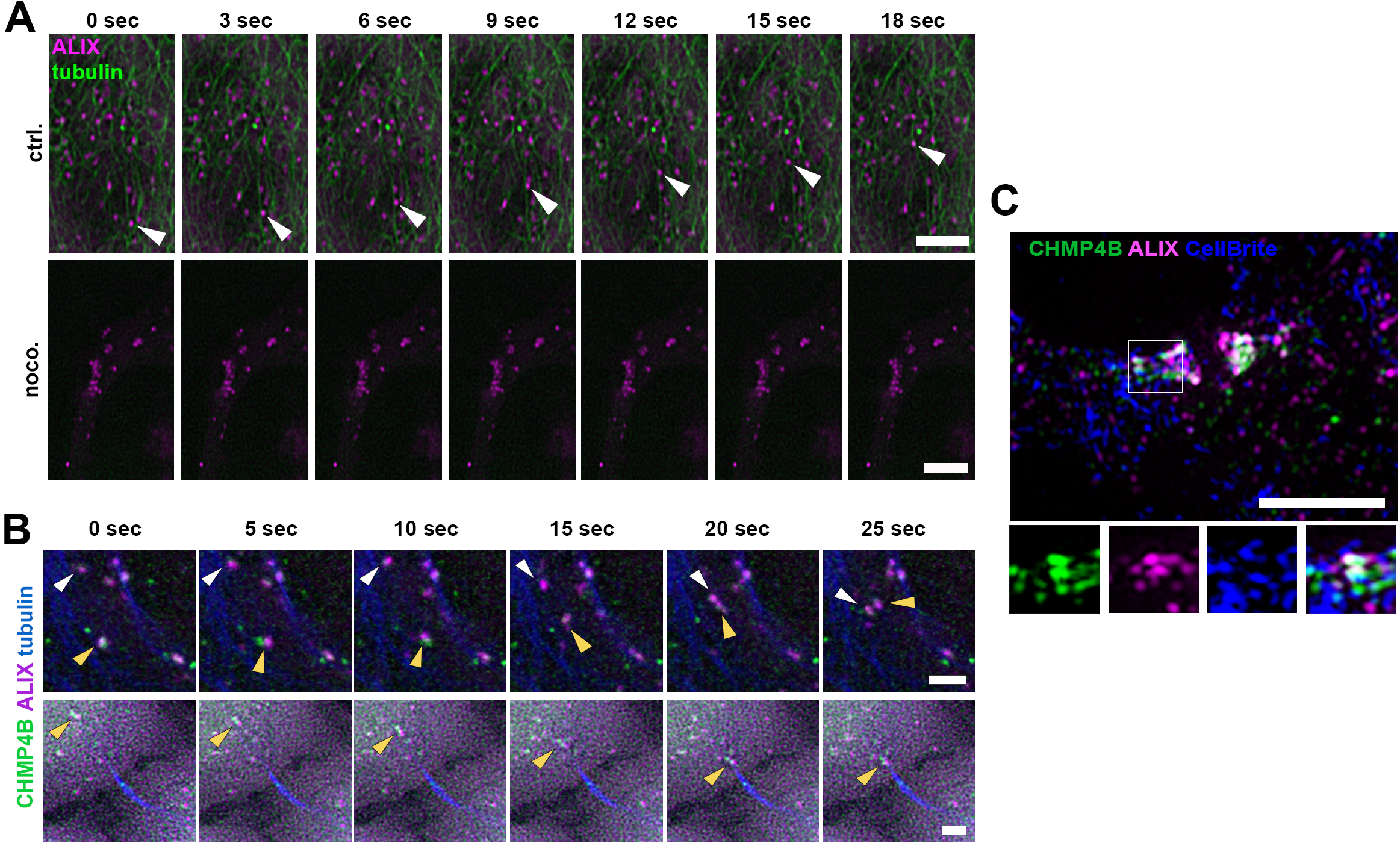
Intracellular transport of ALIX and CHMP4B. **(A)** Selected frames from a time-lapse microscopy video of cells stably expressing ALIX-mCherry upon addition of SiR-tubulin (green) at indicated time points. Top panel: ALIX is associated to vesicles and transported along microtubules. Arrowheads indicate the transport of an individual vesicle. Bottom panel: Nocodazole-mediated microtubule disruption (2h, 60 μM) leads to accumulation and immobilization of ALIX-positive structures. Scale bar = 5 μm. **(B)** Selected frames from time-lapse microscopy videos of cells stably expressing ALIX-mCherry and CHMP4B-GFP upon addition of SiR-tubulin (blue) at indicated time points. Directed co-transport of ALIX and CHMP4B along microtubules in interphase cells (upper panel) and towards the periphery of the intercellular bridge (lower panel). Arrowheads of same color indicate the transport of individual vesicles that are positive for ALIX and CHMP4B. Scale bars = 2 μm. Microtubules are labelled with SiR-tubulin **(A +B)**. **(C)** SIM micrograph of an intercellular bridge of a cytokinetic cell pre-treated for 20h with CellBrite^®^ Steady 650 membrane dye (blue) and stained for ALIX (magenta) and CHMP4B (green). ALIX and CHMP4B signals are associated to membranecontaining vesicles (see also Video 3C and D). Scale bar = 3 μm.

### ALIX- and CHMP4B-positive vesicles can be transported into the intercellular bridge and to the midbody and ALIX is partially co-transported with Rab11

Importantly, in live-cell imaging experiments, ALIX- and CHMP4B-positive vesicles were detected to be transported to the periphery of the intercellular bridge as described above (Video 3D). We also detected partial transport of ALIX- or ALIX- and CHMP4B-positive vesicles into the cytokinetic bridge and visualized recruitment of such vesicles to the midbody (Fig. 4A-B and Videos 4A-C). We furthermore confirmed by super-resolution microscopy of fixed cells a close proximity of endogenous ALIX and CHMP4B on vesicles in the cytokinetic bridge (Fig. 4C and Video 4D).

**Figure 4.**
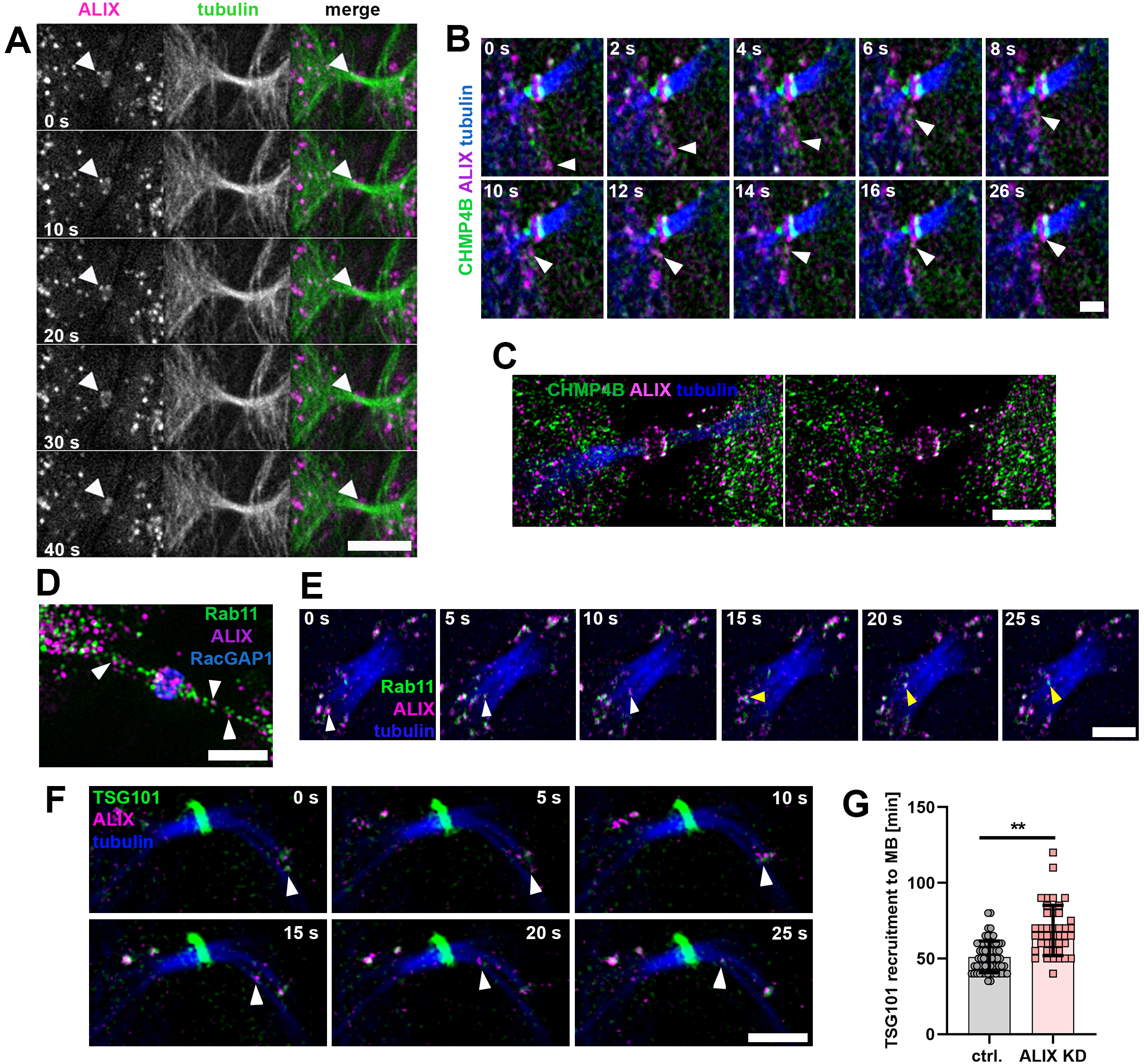
Directed transport of ALIX and CHMP4B to the midbody. **(A)** ALIX transport into the intercellular bridge and recruitment to the midbody and **(B)** ALIX co-transport with CHMP4B into the intercellular bridge. Selected frames at indicated time points from time-lapse imaging of cytokinetic cells stably expressing **(A)** ALIX-mCherry (magenta) or **(B)** CHMP4B-GFP (green) and ALIX-mCherry (magenta) upon addition of SiR-tubulin (green in (A) and blue in (B)). Scale bars = 5 μm (A) and 2 μm (B)). Arrowheads indicate transport of individual vesicles at given time points. **(C)** Colocalization of ALIX with CHMP4B in the cytokinetic bridge. SIM micrographs of fixed cells stained for ALIX (magenta), CHMP4B (green) and tubulin (blue) showing ALIX and CHMP4B co-localization in vesicular structures in the intercellular bridge and at the midbody. Scale bar = 3 μm. **(D)** Colocalization of ALIX with Rab11 in the cytokinetic bridge. SIM micrographs of fixed cells stained for ALIX (magenta), Rab11 (green) and RacGAP1 (blue) show co-localization in vesicular structures in the bridge and at the midbody. Scale bar = 3 μm. **(E)** Co-transport of ALIX and Rab11. Selected frames from a time-lapse microscopy of cytokinetic cells expressing Rab11-GFP (green) and ALIX-mCherry (magenta) with SiR-tubulin (blue). Arrowheads of same color indicate transport of individual vesicles at given time points. Scale bar = 3 μm. **(F)** Co-transport of ALIX and TSG101. Selected frames from a time-lapse microscopy video of cytokinetic cells expressing GFP-TSG101 (green) and ALIX-mCherry (magenta) with SiR-tubulin (blue). Arrowheads indicate transport of an individual vesicle at given time points. Scale bar = 3 μm. **(G)** Scatter plot showing the time interval between bridge formation and appearance of GFP-TSG101 at the midbody in control (ctrl.) cells or upon ALIX knockdown (KD) as indicated (*n* ≥ 26 cells per treatment from four independent experiments; control: 53 ± 1.6 min; ALIX KD: 67.7 ± 3.2 min [±SEM]; P < 0.01).

Interestingly, sometimes ALIX seemed to move in a bi-directional manner in and out of the intercellular bridge or in the periphery of the bridge (Video 4E). Such movement has also been described for Rab11-FIP3 endosomes during early stages of the intercellular bridge of cytokinetic cells (Dambournet et al., 2011; Fremont et al., 2017; Montagnac et al., 2009; Schiel et al., 2011; Takahashi et al., 2011). By SIM of fixed cells, we also found partial co-localization of endogenous ALIX and Rab11 in the proximity of and in the intercellular bridge (Fig. 4D and Video 4F) and FRAP analysis of Rab11 at the midbody showed dynamics resembling those of ALIX and CHMP4B (Suppl. Fig. 2). In addition, live cell imaging confirmed co-transport of ALIX and Rab11, first to the periphery (Video 4G) and finally into the intercellular bridge (Fig. 4E and Video 4H). Besides Rab11-positive vesicles, Rab35 endosomes also promote cytokinesis as described above (Chesneau et al., 2012; Dambournet et al., 2011; Fremont et al., 2017; Klinkert and Echard, 2016; Kouranti et al., 2006). Our FRAP analysis showed different dynamics of Rab35 at the midbody than of Rab11 (Suppl. Fig. 2). Similarly, we could neither detect a high degree of co-transport between ALIX and Rab35 in interphase cells (Video 4I) nor in the intercellular bridge of dividing cells (Video 4J). Our data therefore support the assumption that ALIX transport seems to depend more on Rab11 than on Rab35. Thus, the above data suggest that ALIX and CHMP4B are associated with vesicles and that directional transport of these vesicles occurs, at least partially, via a Rab11-FIP3-mediated process to the intercellular bridge.

### TSG101 associates with vesicles and is transported together with ALIX into the intercellular bridge

During cytokinesis the ESCRT-III component CHMP4B is recruited to the midbody either directly via ALIX or via an ESCRT-I/TSG101-ESCRT-II-CHMP6-dependent mechanism (Carlton et al., 2008; Carlton and Martin-Serrano, 2007; Christ et al., 2016; Morita et al., 2007). Thus, we investigated how the ESCRT-I subunit TSG101 is transported to the midbody. Surprisingly, live cell imaging also revealed colocalization and co-transport between TSG101 and ALIX, both in non-dividing cells (Video 4K) and in the intercellular bridge of dividing cells (Fig. 4F and Video 4L). In addition, knockdown of ALIX resulted in a significantly delayed recruitment of TSG101 to the midbody as compared to its recruitment in control cells (Fig. 4G and Video 4M). This indicates that TSG101 is associated to vesicular structures and that at least a fraction of TSG101 is transported together with ALIX into the intercellular bridge, with this recruitment partially being dependent on ALIX.

### ALIX is co-transported with the kinesin-1 motor protein KIF5B into the intercellular bridge

Finally, we examined by which mechanism ALIX is transported to the midbody. Directed intracellular protein and vesicle transport is mediated via specialized motor proteins that move along the cellular actin cytoskeleton or the microtubule network (Hirokawa, 1998; Sharp et al., 2000). As our data indicated a microtubule-directed transport of ALIX towards and into the cytokinetic bridge, we investigated the role of kinesins (Endow et al., 2010) in this process. This superfamily of microtubule-associated motor proteins includes 14 families of kinesins that mediate an ATP-dependent transport of different cargo along microtubules, and several kinesins have been identified to play a role in mitosis and cytokinesis (Zhu et al., 2005). In particular, we focused on kinesin-1, as this kinesin is a major motor for anterograde transport towards the plus-end of microtubules (Klumpp and Lipowsky, 2005). Furthermore, kinesin-1 is the motor protein for the transport of Rab11-FIP3 vesicles in the cytokinetic bridge (Montagnac et al., 2009). The native conventional kinesin-1 holoenzyme exists as a tetramer consisting of two kinesin heavy chains (KHCs) and two kinesin light chains (KLCs) (DeBoer et al., 2008). In mammals, three genes (*KIF5A, KIF5B*, and *KIF5C*) encode KHC (kinesin-1) isoforms. KIF5A and KIF5C are neuron specific, whereas KIF5B is ubiquitously expressed (Miki et al., 2001; Niclas et al., 1994). Thus, we focused on KIF5B and investigated its role in the transport of ALIX-positive vesicles to the midbody. Live cell imaging revealed a strong co-localization and transport of KIF5B with ALIX-positive vesicles to the periphery of the intercellular bridge (Fig. 5A and Video 5A). High-resolution microscopy confirmed proximity of ALIX and KIF5B (Fig. 5B and Video 5B) as well as of ALIX and the kinesin light chain KLC1 in vesicle-like structures in the cytokinetic bridge (Fig. 5C + Video 5C). These data suggest a co-transport of ALIX and kinesin-1 motor protein KIF5B on vesicles during late stages of cytokinesis.

**Figure 5.**
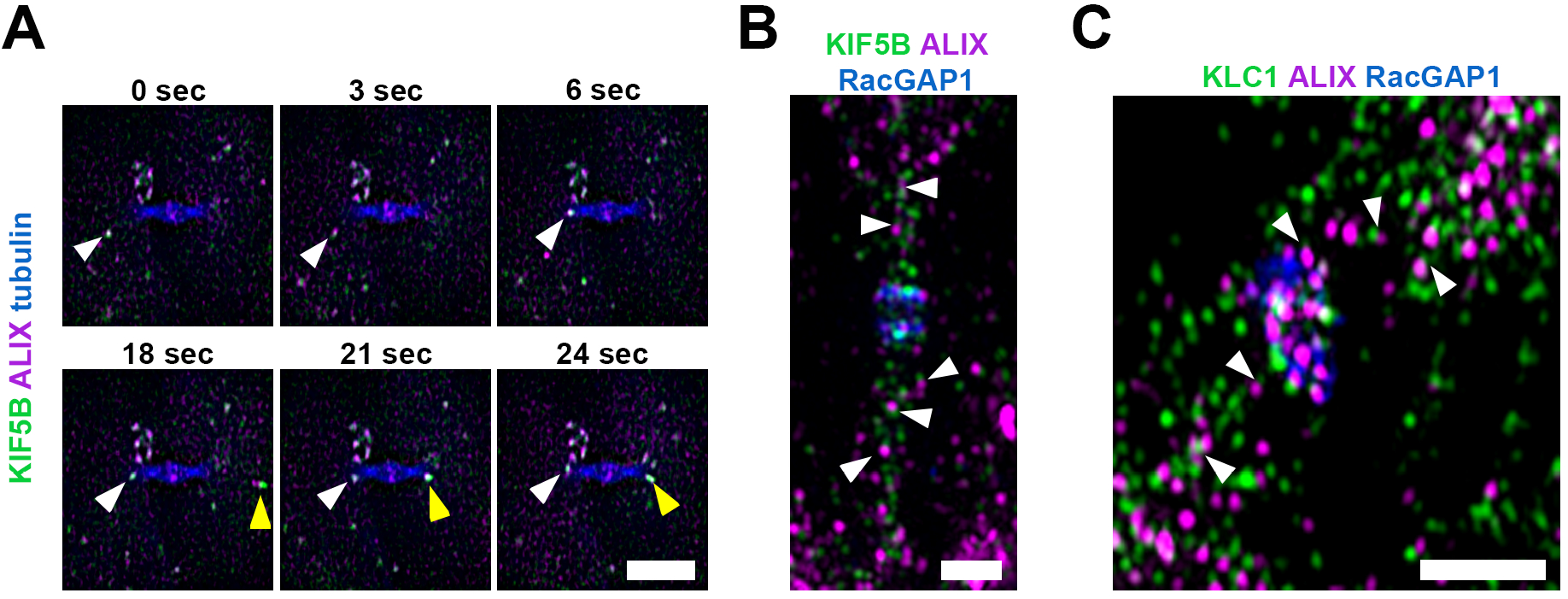
Co-localization and co-transport of ALIX and the kinesin-1 motor KIF5B. **(A)** Co-transport of ALIX (magenta) and KIF5B (green) along microtubules (blue) in an intercellular bridge of a cytokinetic cell. Selected frames from a time-lapse microscopy of cells expressing ALIX-mCherry and mCitrine-KIF5B at indicated time points. Arrowheads of the same color indicate examples of transport of individual vesicular structures that are positive for ALIX and KIF5B. Scale bar = 5 μm. **(B + C)** SIM images of fixed cells stained for ALIX (magenta), RacGAP1 (blue) and KIF5B (green) **(B)** or KLC1 (green) **(C)**. Arrowheads indicate examples of vesicular structures in close proximity positive for ALIX and KIF5B **(B)** or ALIX and KLC1 **(C)**, respectively. Scale bars = 1 μm.

### KIF5B promotes ALIX, CHMP4B and TSG101 recruitment to the midbody and facilitates accurate abscission timing

KIF5B has been shown to promote vesicle transport to the midbody and completion of cytokinesis (Gan et al., 2019; Lawrence et al., 2016; Montagnac et al., 2009). Based on the presence of ALIX and KIF5B co-transport to the intercellular bridge, we asked whether KIF5B plays a role in the transport of ALIX to the midbody and in the completion of cytokinesis. We therefore analyzed the effect of KIF5B knockdown (Fig. 6A) on the abscission time and the recruitment of ALIX to the midbody. Knockdown of KIF5B in HeLa K cells led to a strong delay in the abscission process (Fig. 6B and Suppl. Fig. 3A). This is in accordance with previous reports demonstrating that depletion of KIF5B strongly delays abscission in chondrocytes (Gan et al., 2019). Interestingly, in FRAP as well as in live cell imaging experiments, the dynamics at and recruitment of ALIX to the midbody were significantly delayed upon KIF5B depletion as compared to control cells, respectively (Fig. 6C-D and Video 6A). Importantly, the ALIX protein levels were similar in control HeLa K cells and after KIF5B depletion, as detected by Western blot analysis (Fig. 6A). Moreover, CEP55 protein levels were not reduced in KIF5B-depleted cells (Suppl. Fig. 3B), and KIF5B protein levels were similar in both control and ALIX-depleted HeLa K cells (Suppl. Fig. 3C).

**Figure 6.**
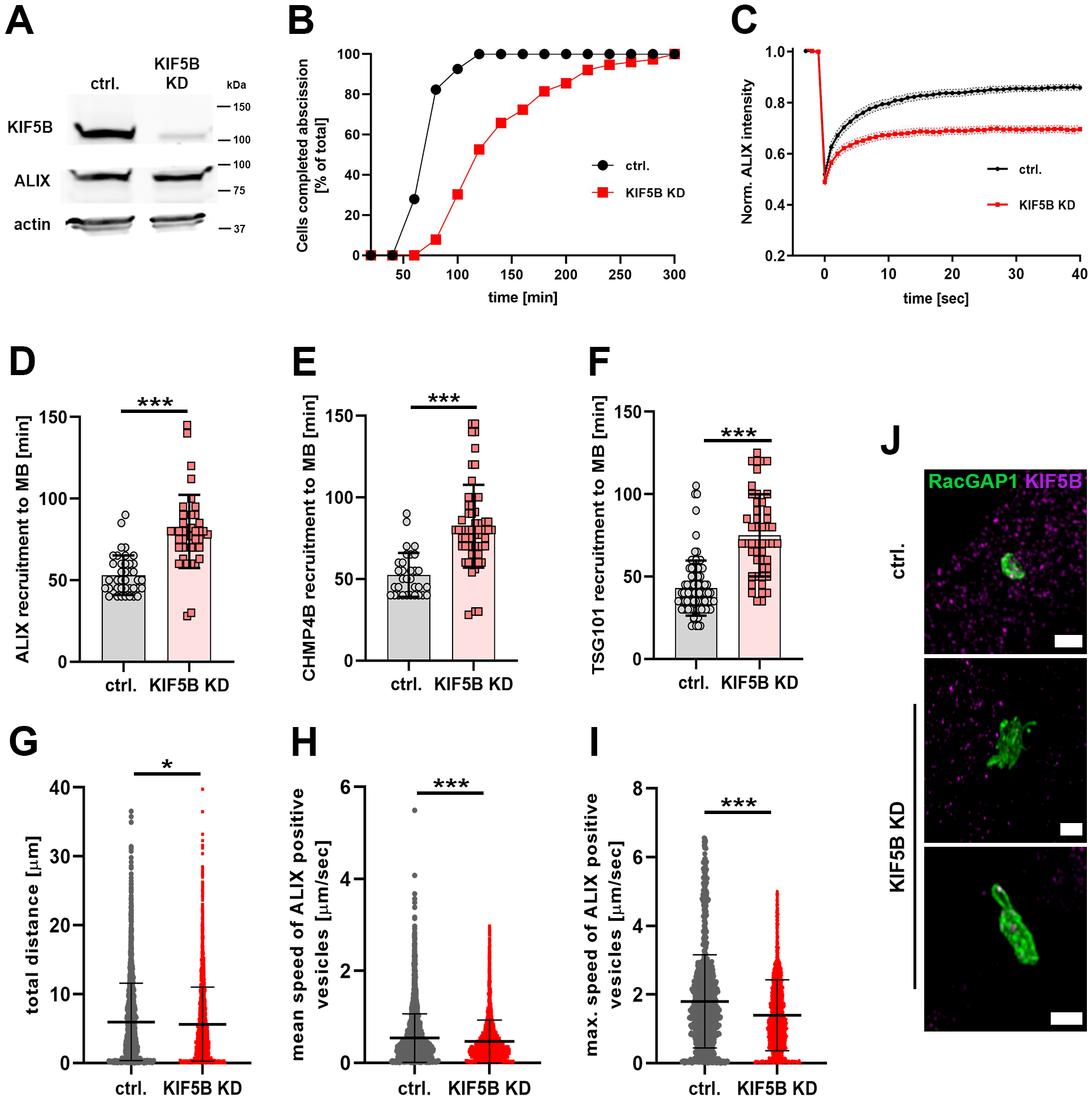
KIF5B promotes ALIX, CHMP4B and TSG101 recruitment to the midbody and accurate abscission timing. **(A)** Western blot showing efficacy of siRNA (12.5 nM) induced KIF5B knockdown (KD) after 2 days of transfection in comparison to constant expression levels of ALIX and actin in both control (ctrl.) and KIF5B siRNA cells. **(B)** Cumulative frequency plot showing the time interval between bridge formation and abscission upon control or KIF5B siRNA treatment as indicated (*n* ≥ 70 cells per treatment from four independent experiments; control: 73.6 ± 2.0 min; KIF5B KD: 141 ± 6.8 min [mean time 50% of cells completed abscission ±SEM]; P < 0.001). **(C)** FRAP analysis of ALIX dynamics at the midbody. Normalized ALIX intensities of control and KIF5B-depleted cells stably expressing ALIX-mCherry are plotted (number of FRAP experiments: ctrl. = 21, KIF5B KD = 16). **(D)** Scatter plot showing the time interval between bridge formation and appearance of ALIX-mCherry at the midbody (MB) in control cells or upon KIF5B siRNA treatment as indicated (*n* ≥ 40 cells per treatment from four independent experiments; control: 53.3 ± 2.3 min; KIF5B KD: 79.4 ± 4.5 min [±SEM]; P < 0.001). **(E)** Scatter plot showing the time interval between bridge formation and appearance of CHMP4B-GFP at the midbody (MB) (*n* ≥ 40 cells per treatment from four independent experiments; control: 52.5 ± 2.4 min; KIF5B KD: 82.3 ± 3.4 min [±SEM]; P < 0.001). **(F)** Scatter plot showing the time interval between bridge formation and appearance of GFP-TSG101 at the midbody (*n* ≥ 40 cells per treatment from four independent experiments; control: 42.7 ± 1.8 min; KIF5B KD: 73.5 ± 3.7 min [±SEM]; P < 0.001). **(G)** Scatter dot plots showing the total distance of ALIX positive vesicles in cytokinetic cells (ctrl. vs. KIF5B KD) during a time period of 7 min (*n* ≥ 30 cells per treatment from four independent experiments; total number of analyzed vesicles ctrl. ≥ 2800 and KIF5B KD ≥ 4700; ctrl.: 5.96 ± 0.10 μm; KIF5B KD: 5.63 ± 0.08 μm [±SEM]; P = 0.0103). **(H)** Scatter dot plots showing the mean speed of ALIX positive vesicles in cytokinetic cells (ctrl. vs. KIF5B KD; *n* ≥ 30 cells per treatment from four independent experiments; total number of analyzed vesicles ctrl. ≥ 4300 and KIF5B KD ≥ 5400; ctrl.: 0.54 ± 0.008 μm/sec; KIF5B KD: 0.47 ± 0.006 μm/sec [±SEM]; P < 0.0001). **(I)** Scatter dot plots showing the maximum speed of ALIX positive vesicles in cytokinetic cells (ctrl. vs. KIF5B KD; *n* ≥ 30 cells per treatment from four independent experiments; total number of analyzed vesicles ctrl. ≥ 1400 and KIF5B KD ≥ 4000; ctrl.: 1.8 ± 0.04 μm/sec; KIF5B KD: 1.4 ± 0.02 μm/sec [±SEM]; P < 0.0001). **(J)** KIF5B depletion affects midbody morphology. Projections of reconstructed 3D SIM data of midbody rings (green, RacGAP1) in fixed control cells or upon KIF5B KD. Depletion of KIF5B (magenta) leads to enlarged and less compact midbody rings. Cells were fixed and stained for RacGAP1 (green) and KIF5B (magenta). Scale bars = 1 μm.

Given the co-transport detected between ALIX and CHMP4B or TSG101 above (Figures 4B and 4F and Videos 4B, 4C and 4M), we also investigated the effect of KIF5B knockdown on the midbody recruitment of CHMP4B or TSG101. Importantly, KIF5B depletion resulted in significantly delayed recruitment of both CHMP4B (Fig. 6E and Video 6B) and TSG101 (Fig. 6F and Video 6C) to the midbody. In line with the delayed midbody recruitment of ALIX, we observed an effect of KIF5B depletion on the motility of ALIX-positive vesicles in cytokinetic cells (Fig. 6G-I). Compared to control cells, KIF5B depletion significantly impaired the total distance travelled by ALIX-positive vesicles (Fig. 6G) as well as the mean and maximum speed of ALIX-positive vesicles (Fig. 6H-I). In addition, similar to the effects induced by ALIX knockdown, the analysis of our 3D SIM data showed structural alterations of the midbody upon KIF5B depletion (Fig. 6J). Visualization of RacGAP1 of the centralspindlin complex revealed considerably enlarged midbodies with irregular shapes and elongated filamentous structures in KIF5B-depleted cells as compared to midbodies in control cells (Fig. 6J and Video 6D). In summary, these data show an important role of KIF5B in mediating cytokinetic abscission by enabling directed transport of proteins required for abscission into the intercellular bridge and to the midbody. In particular, our data provide evidence that the kinesin-1 motor protein KIF5B promotes transport and recruitment of ALIX, CHMP4B and TSG101 to the midbody.

## Discussion

Cytokinesis is a cellular process that demands dramatic morphological reorganizations and is associated with the precise temporal and spatial recruitment of a multitude of different proteins, including ALIX and its associated proteins (Carlton and Martin-Serrano, 2007; Elia et al., 2011; Green et al., 2012; Schoneberg et al., 2017). ALIX is involved in a variety of cellular processes at different cellular localizations (Campsteijn et al., 2016; Sadoul et al., 2018; Vietri et al., 2020). This indicates a highly organized and directed transport of ALIX to conduct a regulated recruitment to the sites of action. However, by which mechanisms cellular transport of ALIX, and in particular, how spatiotemporal recruitment of ALIX to the midbody during cytokinesis occurs, has remained unclear. Here, we present data that, to our knowledge, show a previously uncharacterized directed and kinesin-1-dependent transport of ALIX-positive vesicles along microtubules to the periphery of and into the intercellular bridge, contributing to the accumulation of ALIX at the midbody.

In line with the crucial functional role in cytokinetic abscission, we (Fig. 2B and Suppl. Fig. 1A-B), and others (Addi et al., 2020; Carlton and Martin-Serrano, 2007; Morita et al., 2007), observed a strong delay in abscission upon ALIX depletion. This can be explained by the fact that ALIX recruits further downstream proteins, such as CHMP4B (Carlton et al., 2008; Carlton and Martin-Serrano, 2007; Morita et al., 2007). Accordingly, using live cell imaging, we detected strongly delayed recruitment of CHMP4B to the midbody upon ALIX depletion (Fig. 2C). Furthermore, the delayed abscission process in ALIX-depleted cells was accompanied by elongated intercellular bridges (Fig. 2D). Elongated intercellular bridges upon delayed abscission might be a general phenomenon as it has also been documented as a result of functional inhibition of other cytokinesis-regulating proteins (Carrillo-Garcia et al., 2021). Interestingly, high-resolution microscopy and 3D reconstruction also revealed interference with the structural integrity of the midbody upon ALIX depletion (Fig. 2E). Our findings are in line with Carlton *et al*. who also reported an accumulation of aberrant midbodies after ALIX knockdown (Carlton et al., 2008). Two scenarios seem plausible to us to explain this phenotype. First, ALIX might be needed as a scaffolding and stabilizing component at the midbody. Indeed, a recent publication has shown that ALIX exists at the midbody in complex with syndecan-4, syntenin and CHMP4B (Addi et al., 2020). This complex couples the ESCRT-III machinery to the plasma membrane and therefore stabilizes ESCRT-III at the abscission site and ensures accurate abscission timing (Addi et al., 2020). Alternatively, the absence of ALIX and the accompanied delay of cytokinesis and physical prolongation of intercellular bridges might lead to a deregulation of the mechanical forces affecting the midbody. Certainly, cytokinesis requires precise spatiotemporal regulation of mechanical forces (Andrade et al., 2022; Burton and Taylor, 1997; Carrillo-Garcia et al., 2021; Gupta et al., 2018; Lafaurie-Janvore et al., 2013; Srivastava and Robinson, 2015). Importantly, similar to the ALIX-deficient phenotype, KIF5B depletion also resulted in an altered midbody structure (Fig. 6J). KIF5B functions as a motor protein and KIF5B accumulates in the intercellular bridge adjacent to the midbody at late stages of cytokinesis of chondrocytes (Gan et al., 2019). In summary, the precise spatiotemporal delivery of certain midbody-associated proteins seems to be essential to ensure balanced mechanical forces in the bridge and to maintain the structural integrity of the midbody and midbody.

ALIX is an ESCRT-III-associated protein involved in a variety of ESCRT-III-dependent cellular processes, including cytokinesis. In these processes, ALIX participates in recruiting CHMP4B to the sites of action (Carlton et al., 2008; Christ et al., 2017; Migliano et al., 2022; Morita et al., 2007; Vietri et al., 2020; Zhen et al., 2021). During cytokinesis ALIX promotes recruitment of CHMP4B to the midbody (discussed above) and subsequently CHMP4B polymerizes into helical filaments that form spiral-like structures toward the site of abscission (Addi et al., 2020; Elia et al., 2012; Guizetti and Gerlich, 2012; Guizetti et al., 2011). We detected a high degree of ALIX/CHMP4B co-localization at such spirals at late stages of cytokinetic abscission, identifying that these spirals are also positive for ALIX (Fig. 1C), which is in line with findings by Addi *et al*. (Addi et al., 2020). Generally, it is assumed that the recruitment of CHMP4B to the midbody occurs in dependency of ALIX, but only after its appearance. However, our data show a recruitment or association of ALIX and CHMP4B already outside of the cytokinetic bridge (Fig. 3B) and that subsequently both proteins can be co-transported along microtubules in vesicles to the periphery of and partially into the cytokinetic bridge and then finally accumulate at the midbody (Fig. 4B-C and Videos 3D and 4B). Accordingly, in our experiments we detected an almost identical temporal recruitment of ALIX and CHMP4B to the midbody (Fig. 6D-E). Importantly, at late stages of cytokinesis, ALIX and CHMP4B showed a continuous high degree of co-localization at progressive stages of midbody maturation and spiral formation as detected by super-resolution microscopy (Fig. 1C and Video 1A). Thus, these data propose a new temporal and mechanistic progression in the recruitment of both ALIX and CHMP4B to the midbody.

Live-cell imaging and super-resolution microscopy showed ALIX localized to endosomal vesicles and co-transported together with CHMP4B in interphase cells as well as in cytokinetic cells. In particular, we were able to visualize transport of ALIX/CHMP4B-positive vesicles first to the periphery and subsequently partially into the cytokinetic bridge and to the midbody (Fig. 3B and 4B and Videos 3D and 4B-C). Vesicular transport of ALIX and CHMP4B into the intercellular bridge enables precision in the regulation of their spatial and temporal targeting towards the midbody. An alternative to directional and motor protein-mediated transport is the diffusion of freely accessible ALIX molecules to the sites of action. Indeed, we cannot exclude the possibility that ALIX diffusion might occur in addition to vesicle-associated transport, especially in the cytokinetic bridge and to the midbody. However, a mainly diffusion-driven ALIX distribution would lack complex regulatory mechanism and fast diffusion of vesicle-associated ALIX can physically be excluded. Furthermore, in interphase cells, microtubule dissociation led to aggregation and inhibition of ALIX transport (Fig. 3A), demonstrating the dependency of an intact microtubule network for ALIX transport.

We identified kinesin-1 as an important motor protein for vesicle-associated ALIX, CHMP4B and TSG101 transport to the midbody in HeLa K cells (Figures 5 and 6). KIF5B is the major kinesin-1 motor in non-neuronal mammalian cells and kinesin-1 possesses a crucial role in protein and membrane transport during cytokinesis, in particular the midbody-directed transport of Rab11-FIP3-positive vesicles (Fielding et al., 2005; Montagnac et al., 2009; Wilson et al., 2005). Interestingly, the partial colocalization and co-transport of ALIX and Rab11 that we found particularly in the periphery of the cytokinetic bridge, suggests that a certain fraction of ALIX is co-localized and co-transported on Rab11-FIP3-positive vesicles. On the other hand, it seems likely that ALIX is transported also independently of Rab11, possibly by recruitment to vesicles that lack Rab11. Depletion of KIF5B did not entirely inhibit the recruitment of ALIX to the midbody, but significantly delayed its appearance at the midbody (Fig. 6C-D) and interfered with the dynamics of ALIX-positive vesicles (Fig. 6G-I). However, as we did not obtain a complete depletion of KIF5B, small amounts of KIF5B might be sufficient to mediate a certain ALIX transport. The significantly delayed recruitment of ALIX to the midbody upon KIF5B depletion however suggests that kinesin-1 is an important motor protein for ALIX transport to the midbody during cytokinesis.

In addition to the proximity of ALIX with the kinesin heavy chain KIF5B, we also observed colocalization of ALIX with the kinesin light chain (KLC) KLC1. The kinesin-1 family can consist of four different light chains, which mediate substrate binding (Yip et al., 2016). Additional studies need to be conducted to investigate the extent to which ALIX might associate with the other KLCs and to answer how ALIX or ALIX-positive vesicles are tethered to kinesin-1. To our knowledge, no direct interaction of ALIX with KLCs has yet been documented. Thus, to fully understand mechanisms and regulation underlying the recruitment of ALIX to the midbody, it is important to decipher the detailed interactions between ALIX, kinesin-1, the specific KLCs and further proteins involved.

Our data showed that KIF5B promotes the recruitment of CHMP4B to the midbody (Fig. 6E), which is consistent with its co-transport with ALIX and a similarly delayed recruitment of ALIX (Fig. 6D). Interestingly, KIF5B also promoted accurate recruitment timing of TSG101 to the midbody (Fig. 6F) and ALIX was not only co-localized and co-transported with CHMP4B, but also with TSG101 (Fig. 4F), in the intercellular bridge. Thus, ALIX, CHMP4B and TSG101 are all, at least partially, transported by a KIF5B-dependent mechanism to the midbody.

It seems very likely that beside a KIF5B-mediated ALIX and CHMP4B transport, other mechanisms may occur in parallel or in compensation to reduced KIF5B levels. At least a certain fraction of ALIX and CHMP4B proteins are associated to intracellular vesicles that are transported along microtubules, and this kind of transport depends on motor proteins. Even after KIF5B depletion ALIX can be found associated to motile vesicles (Fig. 6G-I), indicating that either very small amounts of KIF5B are sufficient to maintain ALIX vesicle transport or that their transport can be mediated by other motor proteins. In addition to kinesin-1, other kinesin families have been shown to mediate cargo transport in the intercellular bridge and to the midbody, such as kinesin-2 and kinesin-3 (Keil et al., 2009; Li et al., 2014; Sagona et al., 2010). We assume that besides the recruitment of ALIX and CHMP4B to vesicles, large cytoplasmic amounts of these proteins exist. Thus, in addition to long distance transport along microtubules, local recruitment could also be mediated by cytosolic diffusion. This type of short distance transport could enable a very fast protein recruitment, as we observe during cytokinesis (Fig 1D). Consequently, besides vesicle-mediated transport and recruitment of ALIX and CHMP4B to the midbody, direct recruitment by diffusion would also be feasible. In this case, it would be possible to ensure a high local cytoplasmic concentration of ALIX and CHMP4B by targeted transport of ALIX/CHMP4B-positive vesicles to the periphery of and/or into the intercellular bridge. Indeed, during cytokinesis a large number of ALIX and CHMP4B-positive vesicles were found in the direct periphery of the intercellular bridge (Fig. 3C, 4A, 4C, 4D and Videos 3D and 4A-C). To understand which other mechanisms contribute to the recruitment of ALIX and CHMP4B to the midbody, further experiments are required.

Generally, it is assumed that the midbody is formed by sequential recruitment of the different associated proteins. In contrast to this hypothesis, our findings indicate that ALIX, TSG101 and CHMP4B can be transported in close proximity on identical endosomes on the one hand (Videos 4B-C and 4L), and on the other hand, that knockdown of ALIX delays the recruitment of CHMP4B and TSG101 to the midbody (Fig. 2C and 4G). Thus, it seems possible that recruitment of these proteins might not occur sequentially and separately to the midbody, but that they might already form complexes outside the bridge, which are then transported as a unit to the midbody.

Taken together, our data uncover kinesin-1-mediated directed transport of ALIX, TSG101 and CHMP4B-positive vesicles to and partially into the cytokinetic bridge, which is necessary to ensure normal execution of cytokinesis. Accordingly, depletion of ALIX or KIF5B interferes with the structural integrity of the midbody, the recruitment of midbody-associated proteins, including TSG101 and CHMP4B, and with the abscission process. Further studies are needed to elucidate whether these mechanisms apply in other cell types and species and in ALIX, CHMP4B and TSG101 transport to other cellular targets.

## Materials and methods

### Cell culture, plasmids and transfection

Human HeLa “Kyoto’’ (HeLa K) cells were maintained in DMEM (GIBCO) supplemented with 10% FBS, 100U/ml penicillin, and 100mg/ml streptomycin at 37°C under 5% CO2. Stable cell lines expressing fluorescently labelled ALIX, CHMP4B or TSG101 were previously described (Christ et al., 2016; Radulovic et al., 2018). For transient transfection cells were incubated with mixture of FuGene 6 (Promega) and the plasmid of interest, using a ratio of 3:1 of Fugene 6 to DNA and incubated for 24-48 h. In most cases, cells were transfected in MatTek 3.5 cm dishes using 6 μl of Fugene 6 and 2 μg of DNA. Before cells were used for further analysis the medium was exchanged. The pEGFP-ALIX (Addi et al., 2020), pEGFP-Rab35 and pmCherry-Rab35 (Cauvin et al., 2016; Kuhns et al., 2019) plasmids were a kind gift from Dr. Arnaud Echard, the plasmid expressing Rab11-RFP was kindly provided by Dr. Kay Schink and the plasmid expressing mCitrine-KIF5B was kindly provided by Dr. Eva M. Wenzel. The plasmid expressing ALIX-mCherry (Radulovic et al., 2018), has been described previously.

### siRNA transfections

Silencer Select siRNAs against ALIX (5’-GCAGUGAGGUUGUAAAUGU-3’), KIF5B (5’-CUACAUGAACUUACGGUU-3’) and non-targeting control siRNA (Silencer Select Negative Control No.1 siRNA Cat #4390843) were purchased from Ambion (Thermo Fisher Scientific). Cells were seeded in six-well plates at 30% confluence and transfected with 12.5-25 nM final siRNA concentration using Lipofectamine RNAiMax (Life Technologies) according to the manufacturers’ instructions and cells were then used for experiments 48 or 72 h after knockdown, as defined in the specific Figure legends. Knockdown (KD) cells used in the experiments (ALIX and KIF5B KD) showed ≥80% reduced protein levels compared to control cells, as determined by Western blot analysis.

### Antibodies and other reagents

The following primary antibodies were used for Western blotting (WB) and immunofluorescence (IF): anti-β-actin (Sigma-Aldrich #A5316), anti-ALIX and anti-CHMP4B (Christ et al., 2016), anti-CEP55 (Abnova #H00055165-A01), anti-KIF5B (Abcam #ab151558), anti-GAPDH (Abcam #ab9484) and for IF anti-ALIX (Bio Legend #634502), anti-RacGAP1 (Abcam #ab2270), and anti-tubulin (Sigma-Aldrich #T5168). Secondary antibodies included anti-mouse, anti-rabbit, and anti-goat Alexa Fluor 488 (Jackson ImmunoResearch), Alexa Fluor 555 (Molecular Probes), Alexa Fluor 568 (Molecular Probes),

Alexa Fluor 647 (Jackson ImmunoResearch), and DyLight649 (Jackson ImmunoResearch). Methanol-free 16% paraformaldehyde (PFA) was from Thermo Scientific, SiR700-tubulin from Spirochrome and nocodazole was purchased from Merck.

### Immunoblotting

Cells were washed with ice-cold PBS and lysed in 2× sample buffer (125 nM Tris-HCl, pH 6.8, 4% SDS, 20% glycerol, 200 nM DTT and 0.004% bromophenol blue). Whole-cell lysates were subjected to SDS–PAGE on 4–20% gradient gels (Mini-PROTEAN TGX; Bio-Rad). Proteins were transferred to polyvinylidene difluoride (PVDF) membranes (Trans-Blot^®^Turbo™LF PVDF, Bio-Rad) followed by blocking in 2% BSA and primary antibody incubation in 5% fat-free milk powder in Tris-buffered saline with 0.1% Tween-20 overnight at 4°C. Following three washes with PBS/0.01% Tween-20, the membranes were incubated for 45 min with the fluorescent secondary antibodies IRDye680 or IRDye800 (LI-COR, 926-32212, 926-68073, 1:10,000), washed twice in PBS/0.01% Tween-20 and once in PBS, followed by scanning using an Odyssey infrared scanner (LI-COR). Quantification of immunoblots was performed using ImageJ/FIJI.

### Live-cell microscopy

Live-cell imaging was performed on a DeltaVision OMX V4 microscope equipped with three PCO.edge sCMOS cameras, a solid-state light source and a laser-based autofocus. For long-term live-cell microscopy (12-16 h; analysis of cell abscission, protein recruitment during cytokinesis and length of intercellular bridges) a DeltaVision microscope (Applied Precision) equipped with an Elite TruLight Illumination System, a CoolSNAP HQ2 camera and a 60× Plan Apochromat (1.42 numerical aperture) lens was used. For temperature control during live observation, the microscope stage was kept at 37°C by a temperature-controlled incubation chamber. Cells were imaged in live-cell imaging solution (Invitrogen #A14291DJ) supplemented with 20 mM glucose, 100 U/ml penicillin, and 100 mg/ml streptomycin at 37°C. Time-lapse images (5-10 z-sections, 0.5–2 μm separation) were deconvolved using SoftWoRx software (Applied Precision, GE Healthcare) and processed with FIJI/ImageJ. For visualization of microtubules, cells were pre-treated for 1-3 h with 75 nM SiR-tubulin (Spirochrome). For membrane staining, cells were pre-treated 16-20 h with CellBrite^®^650 according to the manufacturers’ instructions (Biotium). For the analysis of ALIX-vesicle motility, cytokinetic cells were visualized on a OMX 4V microscope (7 min total; 5 sec/frame), processed as described above and analyzed with the TrackMate plug-in in ImageJ (Ershov et al., 2022).

### Structured illumination microscopy (SIM)

For 3D-SIM (structured illumination microscopy) cells were seeded on coverslips and fixed in 4% EM-grade paraformaldehyde for 15 min (Scheffler et al., 2014) and permeabilized with 0.1% Triton X-100 in PBS for 5min. For protein staining, specific primary antibodies were used as defined in the corresponding Figure legends. Coverslips were mounted in ProLong™ Gold or ProLong™ Glass (ThermoFisher). 3D-SIM imaging was performed on a DeltaVision OMX V4 system (Applied Precision) equipped with an Olympus 60x numerical aperture (NA) 1.42 objective, three PCO.edge sCMOS cameras and 405, 488, 568 and 642 nm diode lasers. Z-stacks covering the whole cell were recorded with a Z-spacing of 125 nm. A total of 15 raw images (five phases, three rotations) per plane were collected and reconstructed by using SoftWoRx software (Applied Precision), processed in ImageJ/Fiji (Schindelin et al., 2012) and three-dimensional reconstruction was calculated and visualized by icy imaging software (de Chaumont et al., 2012).

### Fluorescence recovery after photobleaching (FRAP)

For FRAP experiments we used a DeltaVision OMX microscope (Applied Precision, GE Healthcare) with a PlanApo 60×/1.4 NA oil objective. The cells were imaged at a frame rate of 1 frame per second over a period of 40 s (3 s pre-bleach). Spots of the radius *a* = 1.7 μm were bleached with a 488-nm laser at 50% laser intensity for 1 s. Intensity change over time was collected with a 488-nm laser for GFP-conjugated constructs and a 561-nm laser for mCherry or RFP-constructs, respectively. FRAP data were analyzed and recovery curves were plotted after background deduction, normalization of fluorescence intensities (the fluorescence intensities at the last frame before bleaching was set to 100%) and subsequent curve fitting (Carisey et al., 2011).

### Statistical analysis

Statistical analysis was carried out in Graphpad Prism (Graphpad Software). Student’s t-test was used to compare two groups. ANOVA was used to compare multiple groups and Holm–Sidak was used to correct for multiple comparisons. The threshold for significance was set at *P*=0.05. All comparisons made are reported – regardless of significance. Comparisons in the Figures are indicated as n.s.: *P*>0.05, *: *P*<0.05, **: *P*<0.01, ***: *P*<0.001.

## Acknowledgments

We thank Coen Campsteijn, Kay Schink and Eva Wenzel for kindly providing plasmid constructs. We thank the Flow Cytometry Core Facility and the Advanced Light Microscopy Core Facility of Oslo University Hospital for access to instruments. We are grateful to members of the Stenmark Lab for discussions. We would like to thank Arnaud Echard for kindly providing plasmid constructs, discussions, and for critical reading of the manuscript. K.H. acknowledges support from the Norwegian Cancer Society (NCS) (project numbers 163303 and 198147) and the Research Council of Norway (RCN) (project number 288059). S.P. was supported by the grant from RCN (project number 288059). H.S was supported by an Advanced Grant from the European Research Council (project number 788954). This work was partly supported by the Research Council of Norway through its Centres of Excellence funding scheme (project number 262652). The authors declare no competing financial interests.

## Supplementary Figure Legends

**Supplementary Figure 1. Delayed recruitment of CHMP4B to the midbody upon ALIX knockdown.** Selected frames from time-lapse microscopy videos of cells stably expressing CHMP4B-GFP and labelled with SiR-tubulin at indicated time points. **(A)**Recruitment of CHMP4B to the midbody during cytokinesis in control (ctrl.) cells. The upper panel shows SiR-tubulin, visualizing an intercellular bridge, and the bottom panel visualizes the CHMP4B signals. Arrowheads indicate the appearance of CHMP4B at the midbody starting at 50 min after the formation of a stable intercellular bridge. **(B)** Recruitment of CHMP4B to the midbody during cytokinesis in ALIX KD cells. The upper panel shows SiR-tubulin and the bottom panel visualizes the CHMP4B signals. The arrowhead indicates the appearance of CHMP4B at the midbody starting at 120 min after the formation of a stable intercellular bridge. Scale bars = 10 μm.

**Supplementary Figure 2. FRAP analysis of Rab35 and Rab11 dynamics at the midbody during cytokinesis.** Normalized fluorescence intensities from three independent FRAP experiments of cells transiently transfected with either Rab35-GFP (black) or Rab11-GFP (red) are plotted as a function of time (number of FRAP experiments: Rab11 = 39, Rab35 = 26).

**Supplementary Figure 3. Delayed cytokinetic abscission upon KIF5B knockdown. (A)** Selected frames from time-lapse microscopy videos of control cells and KIF5B siRNA-treated cells labelled with SiR-tubulin at indicated time points. In control cells (ctrl., upper panel) the abscission, indicated by cleavage of the intercellular bridge microtubules, takes place 50 min after the formation of a cytokinetic bridge as indicated by an arrowhead. In contrast, abscission in KIF5B-depleted cells (KIF5B KD, bottom panel) is significantly delayed and occurs 120 min after intercellular bridge formation as indicated by an arrowhead. Scale bars = 10 μm. **(B)** Western blots showing efficacy of siRNA (12.5 nM) induced KIF5B knockdown after 2 days of transfection. KIF5B knockdown does not reduce the expression levels of CEP55 and the corresponding GAPDH levels in comparison to control cells (ctrl.). **(C)** Western blots showing efficacy of siRNA (12.5 nM) induced ALIX knockdown (KD) after 2 days of transfection in comparison to constant expression levels of KIF5B and the corresponding actin levels in both control (ctrl.) and ALIX siRNA-induced knockdown cells.

## Video Legends

**Video 1A**

Colocalization of ALIX (magenta) and CHMP4B (green) and formation of spiral like structures at the midbody in fixed cells. Animated projections of 3D reconstructed SIM data from Fig. 1C (middle panel) at different visual angles. Hela K cells were stained for endogenous ALIX and CHMP4B.

**Video 1B**

FRAP analysis of Hela K cells stably expressing ALIX-mCherry. Time-lapse imaging of ALIX-mCherry before and after photobleaching of the ALIX signal in the midbody region. The time between two frames is indicated in seconds. Upon photobleaching (after 3 s) a rapid recovery of the ALIX signal can be detected. For FRAP analysis see Fig. 1D.

**Video 1C**

FRAP analysis of Hela K cells stably expressing ALIX-mCherry. Time-lapse imaging of ALIX-mCherry before and after photobleaching of the ALIX signal in a midbody remnant in post-mitotic cells. The time between two frames is indicated in seconds. Upon photobleaching (after 3 s) no significant recovery of the ALIX signal can be observed. For FRAP analysis see Fig. 1D.

**Video 2A**

Morphology of the midbody ring in untreated Hela K control cells. Animated projections of reconstructed 3D SIM data at different visual angles from Fig. 2E (upper panel). Cells were stained for RacGAP1 (green), CHMP4B (magenta) and ALIX (blue). Control cells show a compact midbody ring with ALIX and CHMP4B accumulation on both sides of the ring structure. In contrast to control cells, ALIX depletion leads to enlarged and less compact midbody rings with filamentous extensions (see also Videos 2B and 2C).

**Video 2B**

Morphology of a midbody ring upon ALIX KD in Hela K cells. Animated projections of reconstructed 3D SIM data at different visual angles from Fig. 2E (middle panel). Cells were stained for RacGAP1 (green), CHMP4B (magenta) and ALIX (blue). Control cells show a compact midbody ring with ALIX and CHMP4B accumulation on both sides of the ring structure. In contrast to control cells (see Video 2A), ALIX depletion leads to enlarged and less compact midbody rings with filamentous extensions (see also Video 2C).

**Video 2C**

Morphology of a midbody ring upon ALIX KD in Hela K cells. Animated projections of reconstructed 3D SIM data at different visual angles from Fig. 2E (bottom panel). Cells were stained for RacGAP1 (green), CHMP4B (magenta) and ALIX (blue). Control cells show a compact midbody ring with ALIX and CHMP4B accumulation on both sides of the ring structure. In contrast to control cells (see Video 2A), ALIX depletion leads to enlarged and less compact midbody rings with filamentous extensions (see also Video 2B).

**Video 3A**

Intracellular transport of ALIX in interphase cells. Time-lapse microscopy videos of cells stably expressing ALIX-mCherry (magenta) and supplemented with SiR-tubulin (green) during the indicated period (in minutes and seconds). In untreated control cells (left) ALIX is associated to vesicles and transported along microtubules (see also corresponding Fig. 3A, upper panels). In nocodazole-treated cells (right) ALIX-positive structures accumulate in perinuclear regions without significant movement (see also corresponding Fig. 3A, lower panels).

**Video 3B**

Intracellular transport of ALIX (magenta) and CHMP4B (green) in cytokinetic cells. Time-lapse microscopy video of cells stably expressing ALIX-mCherry, CHMP4B-GFP and supplemented with SiR-tubulin (blue) during the indicated period (in minutes and seconds). Directed co-transport of ALIX and CHMP4B along microtubules towards the periphery of the intercellular bridge (blue) can be detected. See also corresponding Fig. 3B, lower panel.

**Video 3C**

Intracellular co-transport of CHMP4B (green) and ALIX (magenta) on membrane vesicles. Time-lapse imaging of an interphase cell stably expressing CHMP4B-GFP (green) and ALIX-mCherry (magenta) upon 16h pre-treatment with CellBrite^®^ Steady 650 membrane dye (blue) during the indicated period in minutes and seconds.

**Video 3D**

Co-transport of CHMP4B (green) and ALIX (magenta) on membrane vesicles to the periphery of the intercellular bridge. Time-lapse imaging of a cytokinetic cell stably expressing CHMP4B-GFP (green) and ALIX-mCherry (magenta) upon 16h pre-treatment with CellBrite^®^ Steady 650 membrane dye (blue) during the indicated period in minutes and seconds.

**Video 4A**

Directed transport of ALIX into the intercellular bridge and accumulation at the midbody. Time-lapse imaging of cytokinetic cells stably expressing ALIX-mCherry (magenta) upon addition of SiR-tubulin (green) during the indicated period in minutes and seconds (see also corresponding Fig. 4A).

**Video 4B**

Directed co-transport of ALIX (magenta) and CHMP4B (green) in the periphery and into the intercellular bridge. Time-lapse imaging of a cytokinetic cell stably expressing ALIX-mCherry (magenta) and CHMP4B-GFP (green) upon addition of SiR-tubulin (blue) during the indicated period in minutes and seconds (see also corresponding Fig. 4B).

**Video 4C**

Directed co-transport of ALIX (magenta) and CHMP4B (green) in the periphery and into the intercellular bridge. Time-lapse imaging of three different cytokinetic cells stably expressing ALIX-mCherry (magenta) and CHMP4B-GFP (green) upon addition of SiR-tubulin (blue) during the indicated period in minutes and seconds.

**Video 4D**

Colocalization of ALIX with CHMP4B in the cytokinetic bridge. Animated SIM micrograph of fixed cells stained for ALIX (magenta), CHMP4B (green) and tubulin (blue). Partial co-localization of ALIX and CHMP4B can be detected on vesicular structures in the intercellular bridge and at the midbody (see also Fig. 4C).

**Video 4E**

Bi-directional movement of ALIX in the periphery of the intercellular bridge. Time-lapse microscopy of cells stably expressing ALIX-mCherry during the indicated period (in minutes and seconds).

**Video 4F**

Colocalization of ALIX with Rab11 in the cytokinetic bridge. Animated SIM micrograph of fixed cells stained for ALIX (magenta), RacGAP1 (blue) and Rab11 (green). Partial co-localization of ALIX and Rab11 on vesicular structures in the intercellular bridge (see also Fig. 4D).

**Video 4G**

Co-transport of ALIX and Rab11 in the periphery of the intercellular bridge. Time-lapse microscopy of cells stably expressing ALIX-mCherry (magenta) and transiently transfected with Rab11-GFP (green) upon addition of SiR-tubulin (blue) during the indicated period (in minutes and seconds).

**Video 4H**

Co-transport of ALIX and Rab11 in the periphery and into the intercellular bridge. Time-lapse microscopy of cells stably expressing ALIX-mCherry (magenta) and transiently transfected with Rab11-GFP (green) upon addition of SiR-tubulin (blue) during the indicated period (in seconds; see also corresponding Fig. 4E).

**Video 4I**

Independent transport of ALIX and Rab35 in interphase cells. Live-cell microscopy of cells transiently expressing ALIX-GFP and mCherry-Rab35 upon addition of SiR-tubulin (blue) during the indicated period (in minutes and seconds). ALIX (green) or Rab35 (magenta) vesicles are partially co-localizing in certain endosomal compartments, but no significant co-transport can be observed.

**Video 4J**

ALIX and Rab35 transport during cytokinesis. Live-cell microscopy of cells transiently expressing ALIX-GFP and mCherry-Rab35 upon addition of SiR-tubulin (blue) during the indicated period (in minutes and seconds). ALIX-positive structures (green) are partially transported into the intercellular bridge (blue) whereas no simultaneous transport of Rab35 (magenta) can be observed.

**Video 4K**

Co-transport of ALIX and TSG101 in non-dividing interphase cells. Time-lapse microscopy of cells stably expressing ALIX-mCherry and GFP-TSG101 upon addition of SiR-tubulin (blue) during the indicated period (in minutes and seconds). Vesicles that are positive for ALIX (magenta) and TSG101 (green) are transported along microtubules (blue).

**Video 4L**

Co-transport of ALIX and TSG101 during cytokinesis. Time-lapse microscopy of cells stably expressing ALIX-mCherry and GFP-TSG101 upon addition of SiR-tubulin (blue) during the indicated period (in minutes and seconds). Vesicles that are positive for ALIX (magenta) and TSG101 (green) are transported in the periphery and into the intercellular bridge (blue, see also corresponding Fig. 4F).

**Video 4M**

Delayed recruitment of TSG101 to the midbody upon ALIX knockdown. Time-lapse microscopy videos of cells stably expressing GFP-TSG101 (green) and labelled with SiR-tubulin (blue) during the indicated period (in minutes and seconds). Recruitment of TSG101 to the midbody during cytokinesis in control cells (left) and ALIX KD cells (right). A delayed appearance of TSG101 at the midbody can be observed upon depletion of ALIX (see also corresponding Fig. 4G).

**Video 5A**

Co-transport of ALIX (magenta) and KIF5B (green) to the periphery of the intercellular bridge (blue). Selected frames from a time-lapse microscopy of cells expressing ALIX-mCherry and mCitrine-KIF5B upon addition of SiR-tubulin during the indicated period (in seconds). ALIX- and KIF5B-positive vesicles are transported to the periphery of the intercellular bridge (see also corresponding Fig. 5A).

**Video 5B**

Co-localization of ALIX and KIF5B in the intercellular bridge. Animated SIM micrograph of a fixed cell stained for ALIX (magenta), RacGAP1 (blue) and KIF5B (green). See also corresponding Fig. 5B for indicated sites of proximal localization.

**Video 5C**

Co-localization of ALIX and KLC1 in the intercellular bridge. Animated SIM micrograph of a fixed cell stained for ALIX (magenta), RacGAP1 (blue) and KLC1 (green). See also corresponding Fig. 5C for indicated sites of proximal localization.

**Video 6A**

Delayed recruitment of ALIX to the midbody upon KIF5B knockdown. Time-lapse microscopy videos of cytokinetic cells stably expressing ALIX-mCherry (green) and labelled with SiR-tubulin (blue) during the indicated period (in minutes and seconds). Recruitment of ALIX to the midbody during cytokinesis in control cells (left) and KIF5B KD cells (right). A delayed appearance of ALIX at the midbody can be observed upon depletion of KIF5B (see also corresponding Fig. 6D).

**Video 6B**

Delayed recruitment of CHMP4B to the midbody upon KIF5B knockdown. Time-lapse microscopy videos of cytokinetic cells stably expressing CHMP4B-GFP (green) and labelled with SiR-tubulin (blue) during the indicated period (in minutes and seconds). Recruitment of CHMP4B to the midbody during cytokinesis in control cells (left) and KIF5B KD cells (right). A delayed appearance of CHMP4B at the midbody can be observed upon depletion of KIF5B (see also corresponding Fig. 6E).

**Video 6C**

Delayed recruitment of TSG101 to the midbody upon KIF5B knockdown. Time-lapse microscopy videos of cytokinetic cells stably expressing GFP-TSG101 (green) and labelled with SiR-tubulin (blue) during the indicated period (in minutes and seconds). Recruitment of TSG101 to the midbody during cytokinesis in control cells (left) and KIF5B KD cells (right). A delayed appearance of TSG101 at the midbody can be observed upon depletion of KIF5B (see also corresponding Fig. 6F).

**Video 6D**

Midbody morphology during cytokinesis upon KIF5B knockdown. Animated projections of reconstructed 3D SIM data of the midbody ring (RacGAP1) and KIF5B. Cells were fixed and stained for RacGAP1 (green) and KIF5B (magenta). Control cells show a compact midbody ring (left, see also corresponding Fig. 6J, upper panel). Depletion of KIF5B (magenta) leads to enlarged and less compact midbody rings (middle and right, see also corresponding Fig. 6J, middle and lower panel).

