## Supplementary figures and images for "Vesicle-mediated transport of ALIX and ESCRT-III to the intercellular bridge during cytokinesis"

### Supplemental Figures

**Suppl. Figure 1**

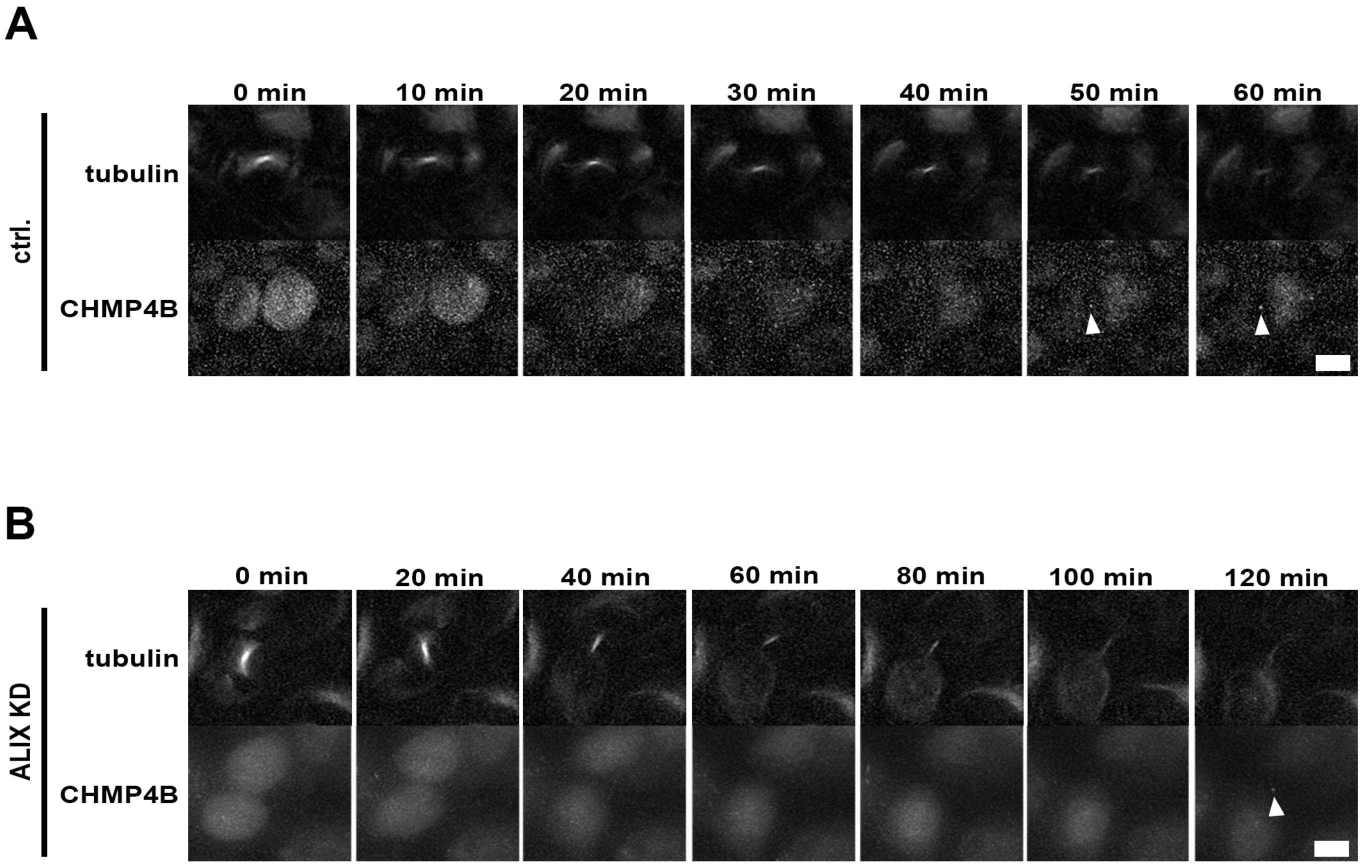

#### Suppl. Figure 2

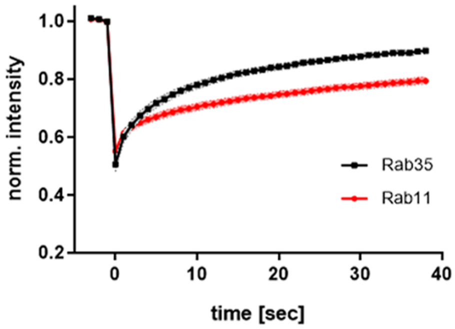

#### Suppl. Figure 3

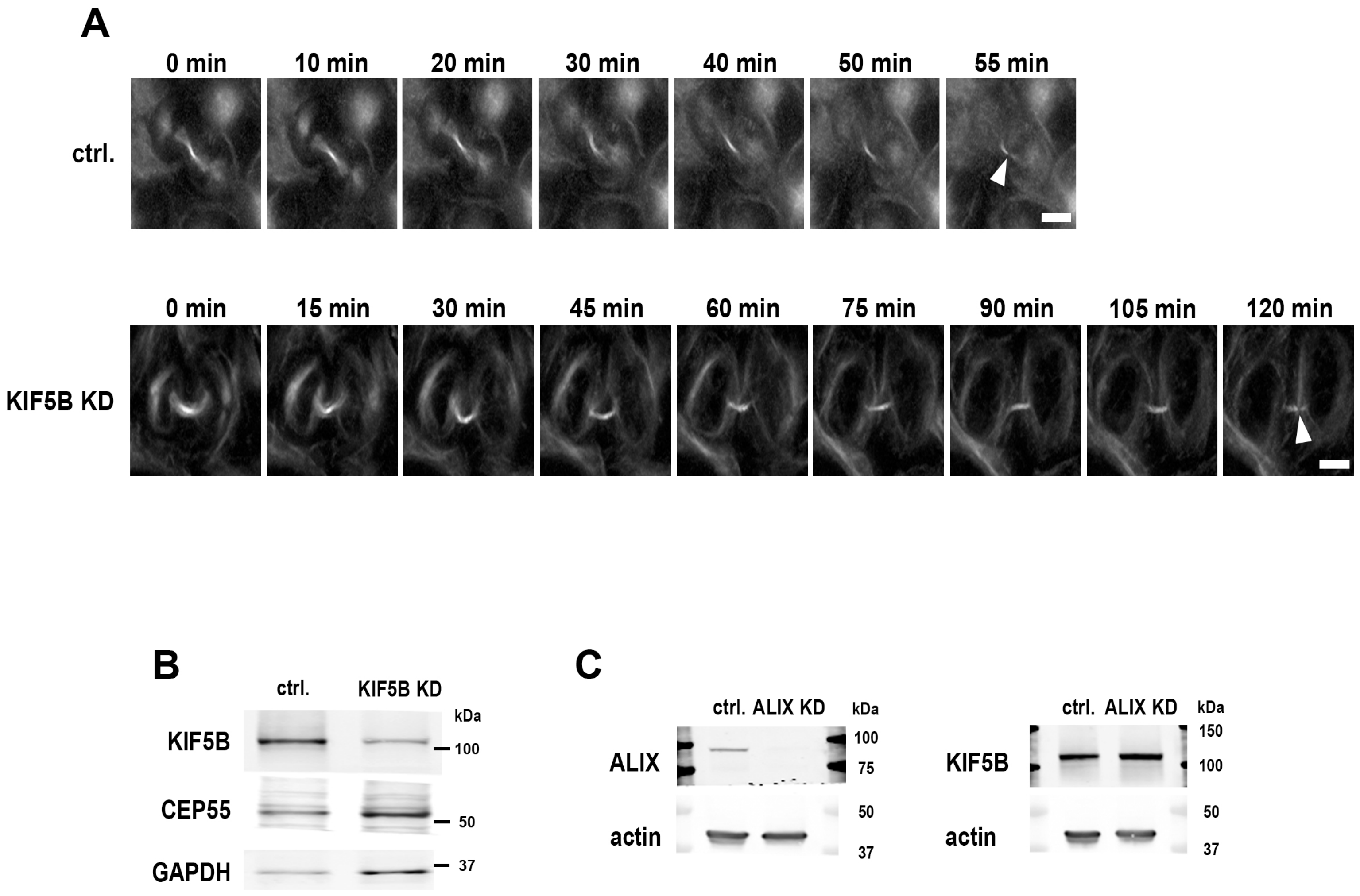
